# AMPK promotes induction of a tumor suppressor FLCN through activation of TFEB independently of mTOR

**DOI:** 10.1101/499921

**Authors:** Caterina Collodet, Marc Foretz, Maria Deak, Laurent Bultot, Sylviane Metairon, Benoit Viollet, Gregory Lefebvre, Frederic Raymond, Alice Parisi, Gabriele Civiletto, Philipp Gut, Patrick Descombes, Kei Sakamoto

## Abstract

AMP-activated protein kinase (AMPK) is a central energy sensor and master regulator of energy homeostasis. AMPK not only elicits acute metabolic responses, but also promotes metabolic reprogramming and adaptions in the long-term through regulation of specific transcription factors/co-activators. We performed a whole-genome transcriptome profiling in wild-type and AMPK-deficient mouse embryonic fibroblasts (MEF) and mouse primary hepatocytes that had been treated with two distinct classes of small-molecule AMPK activators, namely 5-aminoimidazole-4-carboxamide ribonucleotide (AICAR) or 991. This led to the identification of distinct compound-dependent gene expression signatures and to the discovery of several AMPK-regulated genes, including folliculin (*Flcn*), a gene encoding for a tumor suppressor and nutrient sensor. Gene set enrichment and pathway analyses identified the lysosomal pathway and the associated transcription factor EB (TFEB) as key transcriptional mediator responsible for AMPK-dependent gene expression changes. AMPK-induced *Flcn* expression was abolished in TFEB/TFE3 double knockout MEF and the promoter activity of *Flcn* was profoundly reduced when its putative TFEB-binding site was mutated. Mechanistically, we have found that AMPK promotes the dephosphorylation and nuclear localization of TFEB independently of mTOR activity.

Collectively, we identified the AMPK-TFEB-FLCN axis as a potential key regulator for cellular and metabolic homeostasis. Moreover, data from zebrafish with physiologically and pharmacologically activated AMPK confirmed the AMPK-TFEB-FLCN cascade *in vivo*.

## INTRODUCTION

AMP-activated protein kinase (AMPK) is an evolutionary conserved energy sensor which functions to maintain energy homeostasis through coordinating effective metabolic responses to reduced energy availability^1,2^. Low-energy conditions under various pathophysiological settings (*e.g.* nutrient deprivation, physical activity, ischemia), characterized by elevated AMP:ATP or ADP:ATP ratios, trigger AMPK activation. Once activated, AMPK promotes ATP-producing, catabolic pathways and decreases ATP-consuming, anabolic pathways to restore and maintain cellular ATP at a constant level.

AMPK is a heterotrimeric complex composed of a catalytic α-subunit and two regulatory β and γ subunits. Two to three isoforms exist for each subunit (α1 and α2, β1 and β2, and γ1, γ2, and γ3), giving rise to 12 distinct combinations of the heterotrimeric complexes. In general α1, β1, and γ1 appear to be the ubiquitously expressed isoforms of AMPK. There are cell- and tissue-specific distributions of some isoforms (*e.g.* exclusive expression of γ3 in skeletal muscle^3,4^), and they may target AMPK complexes to particular subcellular locations to phosphorylate specific substrates^5,6^. The γ subunits contain four tandem cystathionine β-synthase (CBS) repeats that provide adenine nucleotide binding^7^. AMPK activity increases >100-fold on phosphorylation of a conserved threonine residue within the activation loop (Thr172). Binding of ADP and/or AMP causes conformational changes that favor net Thr172 phosphorylation by the promotion of Thr172 phosphorylation and the inhibition of Thr172 dephosphorylation^8–10^. In addition, the binding of AMP (but not ADP) further promotes AMPK activity by >10-fold by allosteric activation^8^. The major upstream kinase catalyzing AMPKα Thr172 phosphorylation in most mammalian cells/tissues, including skeletal muscle and liver, is the tumor suppressor kinase LKB1^11^. In some cell types, Thr172 can be phosphorylated in a Ca^2+^-mediated process catalyzed by Ca^2+^/calmodulin-dependent protein kinase kinases^12^.

AMPK is considered an attractive therapeutic target for metabolic disorders since AMPK activation brings about metabolic responses anticipated to counteract the metabolic abnormalities associated with obesity, insulin resistance, and type 2 diabetes^13,14^. Indeed several compounds, which can be divided into three categories^12^, have been reported to activate AMPK and elicit metabolic effects in cellular and pre-clinical studies. The first class comprises indirect activators which, through inhibition of mitochondrial respiration and eventual suppression of ATP synthesis, increase cellular AMP:ATP or ADP:ATP ratios (*e.g.* metformin, resveratrol)^15^. The second class includes pro-drugs converted to AMP analogs inside the cells, and the most well characterized and commonly used compound is 5-aminoimidazole-4-carboxamide-1-β-D-ribofuranoside (AICAR)^16,17^. The allosteric activators, binding to a site located between the α-subunit kinase domain and the β-subunit carbohydrate binding module (CBM) termed “allosteric drug and metabolite (ADaM)” site, constitute the third class. The first compound identified through a high-throughput screen^18^ as an allosteric AMPK activator is A769662, a thienopyridone that preferentially activates β1-containing complexes^19^. More recently 991 was developed, a cyclic benzimidazole which binds AMPK 10 times tighter than A769662 in cell-free assays^20^ and activates both β1- and β2-containing complexes (with a higher affinity to the β1 complexes)^4,21^. It has been reported that A769662 causes several off-target effects, for example in isolated mouse skeletal muscle when used at high concentration^22^ and also in other cells/tissues^23,24^ and was reported to have poor oral availability^18^. Notably, an emerging new generation of ADaM site-binding compounds, including MK-8722 and PF-739, have been shown to be effective in reversing elevated blood glucose concentrations in rodents and non-human primates through activation of AMPK *in vivo*^25,26^.

It is well established that AMPK elicits a plethora of acute metabolic responses through phosphorylation of serine residues surrounded by the well characterized recognition motif^27^. There has been much effort into the identification of AMPK substrates, and several targeted and untargeted proteomics studies have been performed^28–32^, which led to a mechanistic understanding of AMPK-mediated metabolic responses and the discovery of new roles for AMPK beyond conventional metabolic regulation (*e.g.* cell cycle, autophagy). Growing evidence suggests that in the long-term AMPK promotes metabolic reprogramming via effects on gene expression at least partly through regulation of specific transcription factors and transcriptional co-activators^1,12^.

In the current study we initially performed a comprehensive transcriptomic profiling in wild-type (WT) and AMPK-deficient (AMPKα1/α2 double knockout (AMPK KO)) mouse embryonic fibroblasts (MEF) and primary hepatocytes treated with AICAR and 991, which led to the identification of distinct compound-dependent gene expression signatures and to the discovery of several AMPK-regulated genes. Gene set enrichment and pathway analyses prompted us to hypothesize that the transcription factor EB (TFEB) is a potential key transcription factor responsible for AMPK-mediated gene expression. We found that expression of *Flcn*, a gene which encodes the tumor suppressor folliculin (FLCN), was abolished in both AMPK KO and TFEB/TFE3 double KO MEF and the promoter activity of *Flcn* was profoundly reduced when the putative TFEB-binding site was mutated. Finally, we found that AMPK activates TFEB through promotion of dephosphorylation and nuclear translocation independently of mTOR signaling.

## RESULTS

### Whole-genome transcriptome profiling revealed distinct gene expression profiles in response to AMPK activators in MEF and primary hepatocytes

To identify genes and pathways regulated in an AMPK-dependent mechanism, we performed a whole-genome transcriptome profiling using microarray technology (Affymetrix Mouse GeneChips). Taking cell type-specific roles and isoform-/compound-selective responses of AMPK into account, we used two different cell models and genotypes, namely AMPK WT and AMPK KO in both MEF and mouse primary hepatocytes, and treated them with two different AMPK activators (991 and AICAR) known to target distinct regulatory sites/mechanisms^27^. MEF and mouse primary hepatocytes were treated with vehicle, 991 or AICAR at the indicated concentrations for 4 hours. Following treatment, one set of the samples was subjected to transcriptome profiling and the other set was used for immunoblotting to assess the effect of compounds on AMPK activity. We initially confirmed by immunoblot analysis that treatment of WT MEF with 991 (10 μM) or AICAR (2 mM) resulted in an increase in phosphorylation of AMPK (Thr172) and its *bona fide* substrates acetyl CoA carboxylase (ACC) (Ser79) and raptor (Ser792) (Fig. 1A). Similar results were obtained when WT mouse primary hepatocytes were treated with 991 (3 μM) or AICAR (300 μM) (Fig. 1B). Notably, although treatment with 991 or AICAR resulted in a comparable elevation of ACC phosphorylation in both MEF and hepatocytes, 991-induced raptor phosphorylation was higher in MEF but lower in hepatocytes compared to AICAR (Fig. 1A, B). As previously demonstrated^33–35^, there was no detectable phosphorylation of AMPK, ACC and raptor in vehicle- and 991-/AICAR-treated AMPK KO MEF and hepatocytes (Fig. 1A, B). Taken together, we validated a complete ablation of AMPK activity in both AMPK KO MEF and hepatocytes, and observed differential responses in commonly used surrogate markers for cellular AMPK activity (i.e. phosphorylation of ACC and raptor) following the treatment with 991 or AICAR.

**Figure 1.**
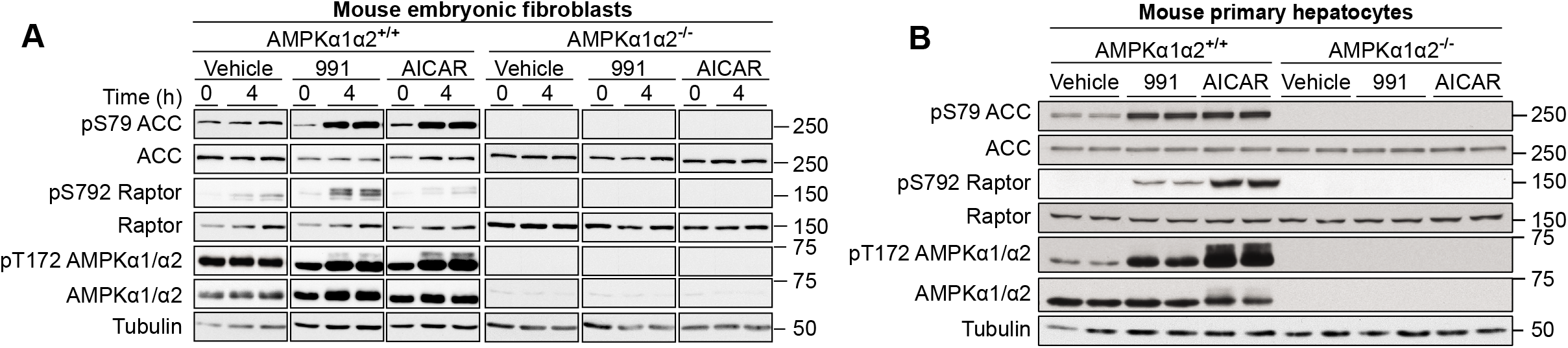
AMPK-activation in samples used for transcriptomic analyses. (**A**) Mouse embryonic fibroblasts control (AMPKα1α2^+/+^) or knockout (AMPKα1α2^-/-^) were treated with vehicle (DMSO), 2 mM AICAR or 10 μM 991 for 4 h. Cell lysates (20 μg) were subjected to western blot analysis with the indicated antibodies and a representative blot of n=3 is shown. (**B**) Primary hepatocytes were isolated from AMPKα1/α2 liver-specific knockout (AMPKα1α2-/-) mice and control AMPKα1^lox/lox^ α2^lox/lox^ mice littermates (AMPKα1α2^+/+^). Plated cells were treated for 4 h with vehicle (DMSO), 3 μM 991 or 300 μM AICAR. Laemmli extracts (20 μg) were subjected to western blot analysis using the indicated antibodies (n=2).

We next performed a whole transcriptome profiling using Affymetrix MOE430 arrays. A principal component analysis (PCA) was conducted for initial evaluation of data quality and assessment of the effect of genotype and treatment on the transcriptome (Supplementary Fig. 1A, B). In MEF the PCA clarified that genotype was the primary factor responsible for variance, followed by treatment, and that globally AICAR caused a greater transcriptional response compared to that induced by 991. It also implied that 991 induced a rather limited, but potentially more specific transcriptional responses compared to AICAR. In contrast, in primary hepatocytes, treatment was the main cause of variance, followed by a lower variance explained by genotype. AICAR consistently produced a much greater response than that driven by 991. These observations were well corroborated by the hierarchical clustering (Fig. 2A, B).

**Figure 2.**
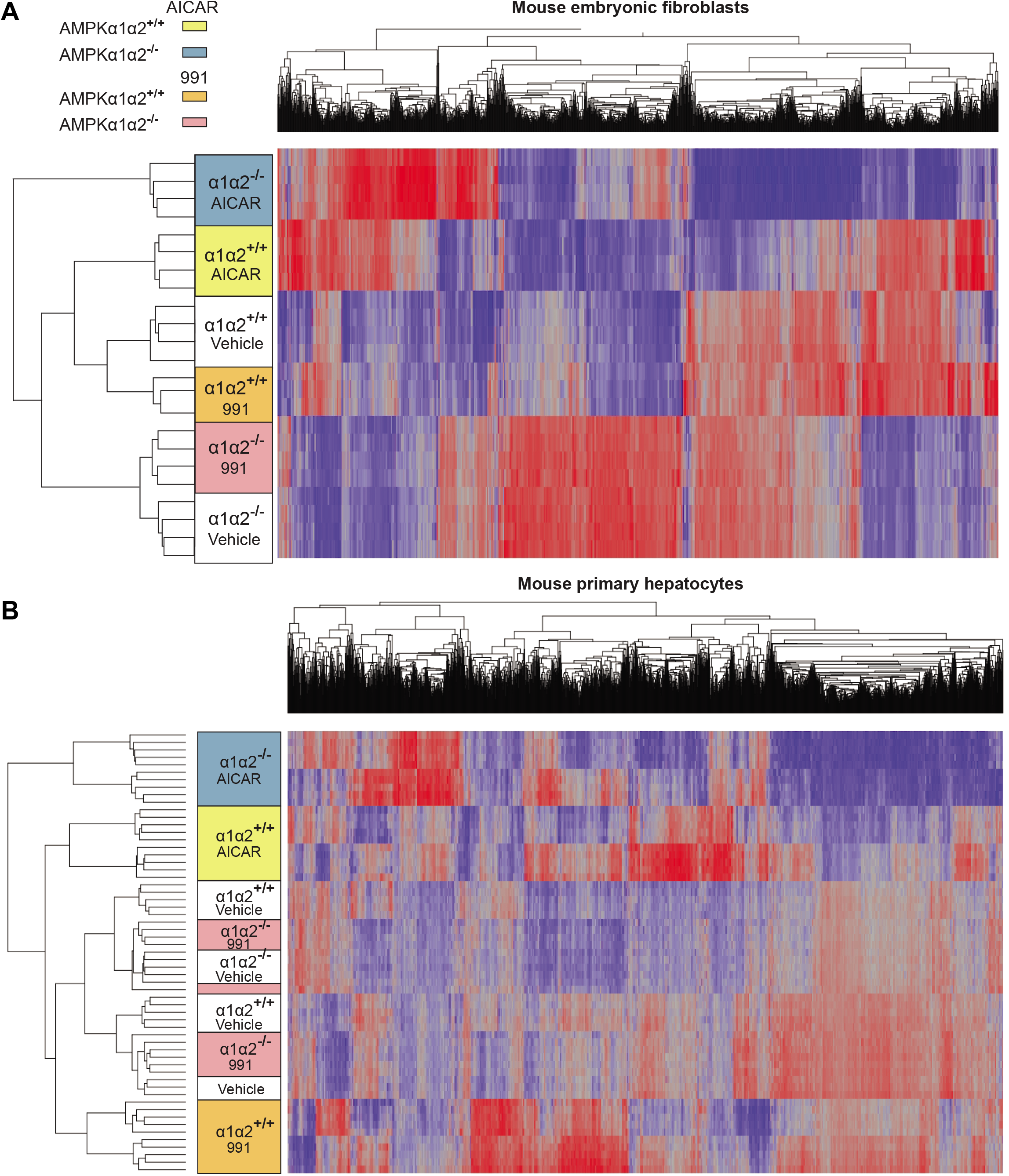
Hierarchical clustering of the transcriptomic data. Overview of the hierarchical cluster analyses in mouse embryonic fibroblasts (**A**) and mouse primary hepatocytes (**B**). Mean-centered gene expression ratios are depicted by a log2 pseudo color scale, red indicates that the gene is overexpressed compared to the mean value whereas blue specifies that the gene is less expressed. Data were analyzed by two-way ANOVA with the factors of genetic background, treatment and interaction. Only selected genes, having a false discover rate (FDR) *P* < 0.0001 for the interaction, are shown.

### Identification and validation of AMPK-dependent genes and pathways

To clarify genes specifically regulated following AMPK activation, we first selected genes by pairwise differential analysis of MEF and primary hepatocytes treated with AICAR or 991 as compared to vehicle. *P*-values were corrected for multiple testing using the false-discovery rate (FDR) method of Benjamini and Hochberg^36^ and we applied a conservative significance threshold of 5% FDR associated with a fold change value of 1.3 or more, given that moderate fold changes were observed. Following 991 treatment, the vast majority of differentially expressed transcripts (>92%) required a functional AMPK, with 184 out of 199 for MEF and 670 out of 684 for primary hepatocyte transcripts regulated in an AMPK-dependent fashion, respectively (Fig. 3A, B and Supplementary Table 1). This observation confirms the nearly exclusive specificity of 991 for targeting AMPK in MEF and hepatocytes consistent with the *in vitro* (cell-free) observation in our previous study^4^. In contrast, AICAR induced a much greater transcriptional response with a majority (~50%) of the transcripts differentially regulated in the absence of AMPK (1026 out of 2053 in MEF and 754 out of 1718 in primary hepatocytes, respectively) (Fig. 3A, B). Altogether, these findings suggest that 991 elicits much more AMPK-specific transcriptional responses compared to AICAR, an observation which is corroborated by the results of PCA (Supplementary Fig. 1) and hierarchical clustering (Fig. 2A, B).

**Figure 3.**
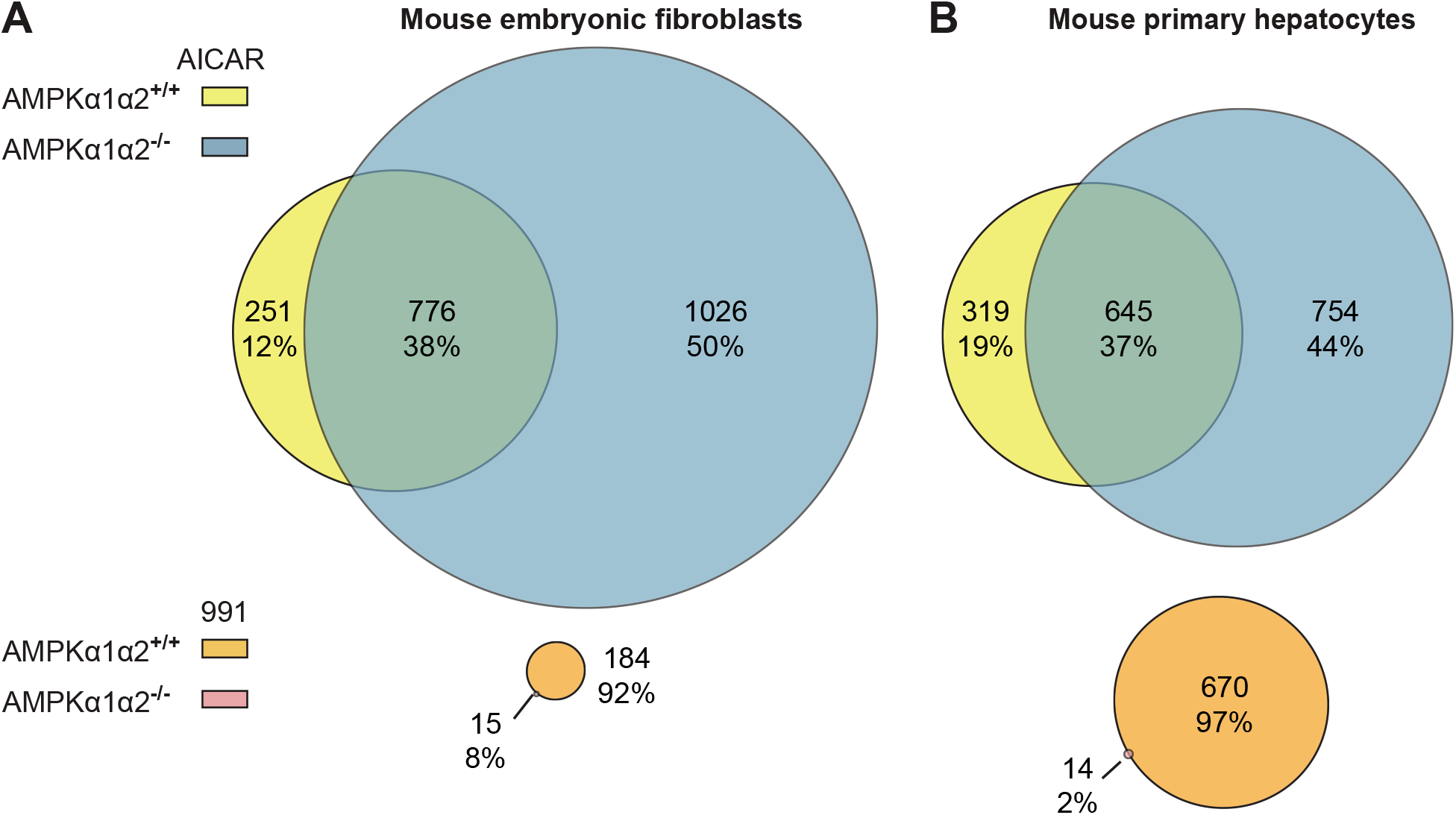
Transcriptomic data analysis of the AMPK-activation response following treatment with 991 and AICAR. Venn diagrams showing the transcriptome profiling specificity of 991 and AICAR in mouse embryonic fibroblasts (**A**) and mouse primary hepatocytes (**B**). Two-way analysis of variance (ANOVA) with Benjamini & Hochberg multiple testing correction was applied to discriminate 991 versus control and AICAR versus control conditions. The moderated *P*-value was set at 0.05 for the interaction between the genetic background and treatment, as well as for the pairwise comparisons, and a fold-change cutoff of 1.3 was applied. The color corresponds to the two treatment conditions and two cellular models as indicated in the legend. The numbers correspond to the numbers of transcripts altered and the percentage contribution to the total number of transcripts identified for each treatment.

To illuminate the pathways regulated by AMPK, we performed a gene ontology analysis on the genes differentially expressed upon 991 treatment in primary hepatocytes and MEF using the Database for Annotation, Visualization and Integrated Discovery (DAVID) program^37,38^ (Fig. 4 A, B). This analysis revealed a commonly shared signature of biological/metabolic pathways (*e.g.* lysosomes) observed in both cell types, as well as additional cell type-specific pathways such as ErbB signaling, sphingolipid metabolism and adipocytokine signaling for primary hepatocytes, and steroid biosynthesis and biosynthesis of antibiotics for MEF, respectively (Fig. 4A, B). In order to validate the microarray data, we performed quantitative real-time PCR (qPCR) analyses on several genes that are known to be involved in lipid/cholesterol signaling and metabolism (*Acss2, Crebrf, Hmgcr, Ldlr, Lpin1, Msmo1, Pde4b*), glucose transport (*Txnip*), immunity (*Ifit1, Tollip*), and cell growth/cancer (*Flcn, Fnip1, Mnt*) (Fig. 4C, D). Overall, we observed a strong correlation between the microarray and qPCR data. Additionally, we also found cell type-/compound-specific responses. For example, AICAR had a more profound effect on gene expression compared to 991 in MEF (*Acss2, Flcn, Ldlr, Lpin1, Mnt, Tollip, Txnip*), but this was not the case in primary hepatocytes (Fig. 4C, D). Interestingly, while AICAR exhibited significant and greater effects on expression of *Flcn* and *Fnip1* in MEF compared to 991, it had no significant effect in primary hepatocytes. In addition, AICAR elicited an AMPK-dependent effect on *Hmgcr* and *Lpin1* expression in MEF, which was absent in primary hepatocytes. Collectively, we identified AMPK-dependent genes (involved in metabolism, immunity and cell growth) that demonstrate cell type- and compound-specific responses in MEF and mouse primary hepatocytes.

**Figure 4.**
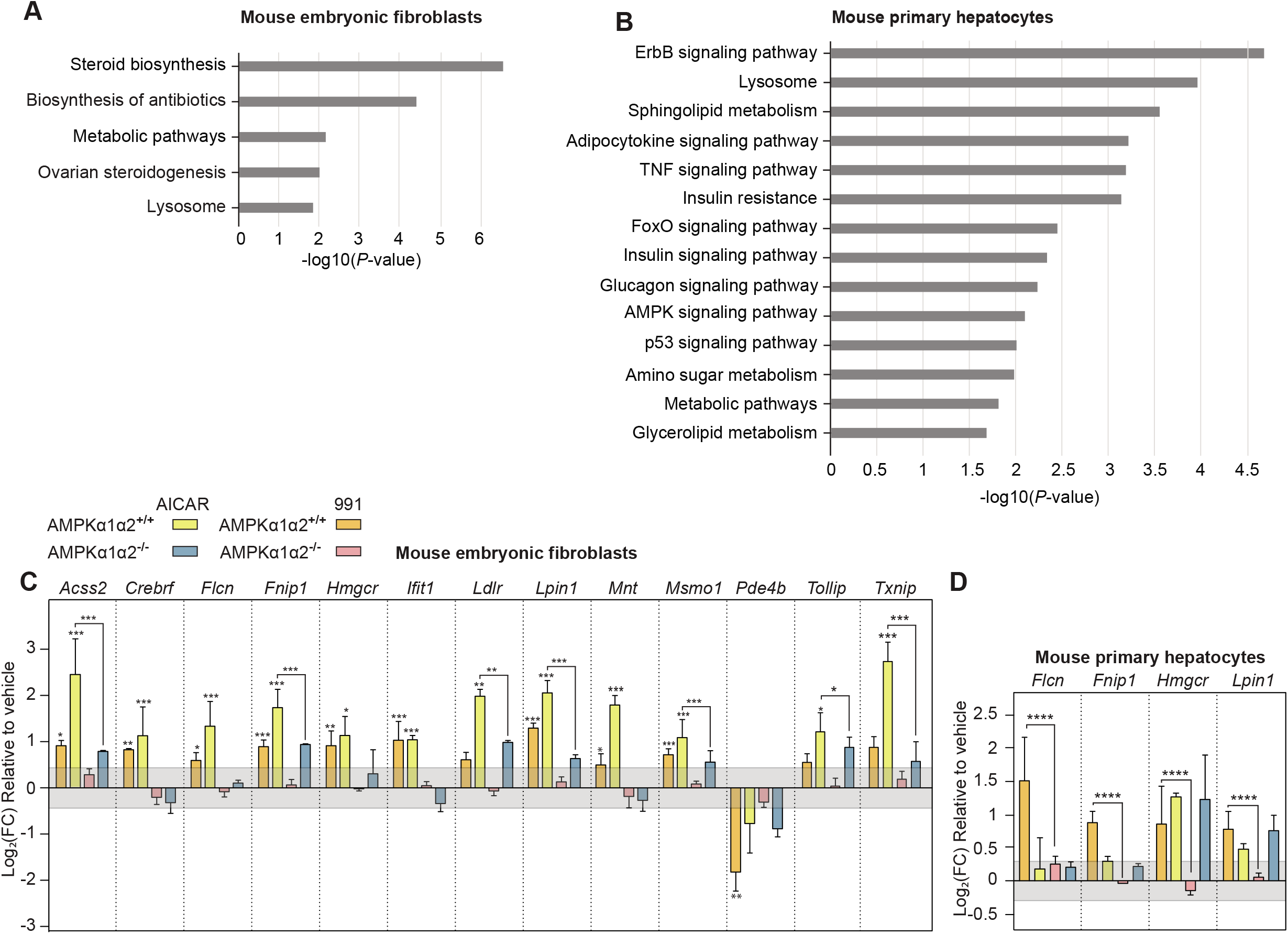
Identification of pathways and genes modulated by AMPK. Gene enrichment analysis of the 991-responsive signature in mouse embryonic fibroblasts (**A**) and mouse primary hepatocytes (**B**). DAVID was used to explore the gene ontology terms associated to the AMPK-regulated genes. The bars represent the negative log10(*P*-value) of enriched terms, indicating the significance of association between the gene list and an indicated ontology term. Relative mRNA levels of the indicated genes, in mouse embryonic fibroblasts (**C**) and primary hepatocytes (**D**). AMPKα1α2^+/+^ or AMPKα1α2^-/-^ MEF were stimulated with 10 μM 991 or 2 mM AICAR for 4 h, and AMPKα1α2^+/+^ or AMPKα1α2^-/-^ hepatocytes were treated with 3 μM 991 or 300 μM AICAR for 4 h. The color corresponds to the two treatment conditions and two cellular models as indicated in the key. Three normalization genes were used (Acyl-CoA synthetase short-chain 2 *Acss2*, β2 microglobulin *B2m*, Peptidylprolyl isomerase-a *Ppia*), and their stability was assessed using GeNorm (M value < 0.6). Values are represented as log2 fold-change of the mean ± SD (n=9). The grey shaded area indicates the log2 fold-change threshold of ±0.37. For the analysis, a two-way ANOVA with interaction was fit to log-transformed data, \**P* < 0.05, \*\**P* < 0.01, \*\*\**P* < 0.001.

### 991-/AICAR-stimulated *Flcn* expression is AMPK-TFEB/TFE3 dependent

To shed light on the mechanism by which AMPK modulates expression of specific genes, we next conducted an *in silico* analysis to identify potential transcription factors responsible for the AMPK-dependent responses upon compound treatment. To this end, we performed a search for correlation in expression between the 991-responsive transcripts identified in MEF (184 targets) and hepatocytes (670 targets), and the expression of potential upstream regulators using the Ingenuity Pathway Analysis (IPA) tool^39^ (Table 1). The IPA revealed that there were around 10 candidates and only two of these, sterol regulatory element binding protein 1 (SREBP1) and TFEB, were identified in both MEF and hepatocytes (Table 1). SREBP1 is a master transcriptional regulator of lipid synthesis and its activity is known to be regulated by AMPK-mediated phosphorylation^40^. TFEB and TFE3 are members of the micropthalmia (MiT/TFE) family of HLH-leucine zipper transcription factors that play an important role in the control of cell and organismal homeostasis through regulating lysosomal biogenesis and autophagy^41^. TFEB is also known to be indirectly regulated by AMPK in the control of lineage specification in embryoid bodies^42^. Interestingly, it was recently reported that TFEB/TFE3 regulate energy metabolism, although the underlying mechanism remains elusive^43,44^. Among the genes regulated specifically in an AMPK-dependent manner (Fig. 4C, D), we focused on *Flcn*, which encodes the tumor suppressor folliculin (FLCN). FLCN and its binding partner FLCN-interacting protein (FNIP) are known to interact with AMPK and this FLCN-FNIP-AMPK interaction/complex has been proposed to control various metabolic functions^45–47^. We first determined the kinetics of the effects of 991 and AICAR on the expression of *Flcn* in time-course experiment over 24 hours, followed by qPCR analysis. We observed that 991 and AICAR induced significant and prolonged expression of *Flcn* throughout the time points measured in an AMPK-dependent manner compared to the vehicle control, except the 24 hour time point where AICAR caused an increase in *Flcn* expression in AMPK KO MEF (Fig. 5A). We confirmed that the increased levels of *Flcn* mRNA were due at least partly to enhanced gene transcription, since pre-mRNA levels of *Flcn* were also significantly elevated in response to 991 and AICAR treatment (Supplementary Fig. 2A, B). Moreover, we confirmed that the elevated levels of *Flcn* transcripts were translated into an increase in FLCN protein levels in a time- and AMPK-dependent manner in response to 991 or AICAR treatment (Fig. 5B). We next wanted to determine if TFEB is required for mediating AMPK-induced expression of FLCN at both mRNA and protein levels. Since TFEB and TFE3 have been proposed to be at least partially redundant^41^, TFEB/TFE3 double KO MEF were employed. As anticipated 991 or AICAR significantly increased expression of *Flcn* at both mRNA and protein levels in WT, but not in TFEB/TFE3 double KO MEF (Fig. 5C, D). We hypothesized that TFEB regulates *Flcn* expression through enhancing its promoter activity, especially based on the fact that the *Flcn* promoter contains a putative TFEB binding site, which is conserved between humans and mice. To test this hypothesis, we cloned and placed fragments of various length (8000, 1200, and 100 bp) of the mouse *Flcn* promoter region upstream of the luciferase gene and transfected the resulting reporter construct individually into WT or AMPK KO MEF. 991 increased the promoter activity of all three constructs when introduced into WT MEF, while in contrast no or only a marginal increase was seen when they were introduced into AMPK KO MEF (Supplementary Fig. 2C). Notably, when the putative TFEB binding site was mutated in the 1200 and 100 bp fragments and the corresponding reporter constructs introduced in WT MEF, 991-induced promoter activity was significantly reduced (Supplementary Fig. 2D). Taken together, these results demonstrate that AMPK promotes *Flcn* transcription through activation of TFEB/TFE3.

**Figure 5.**
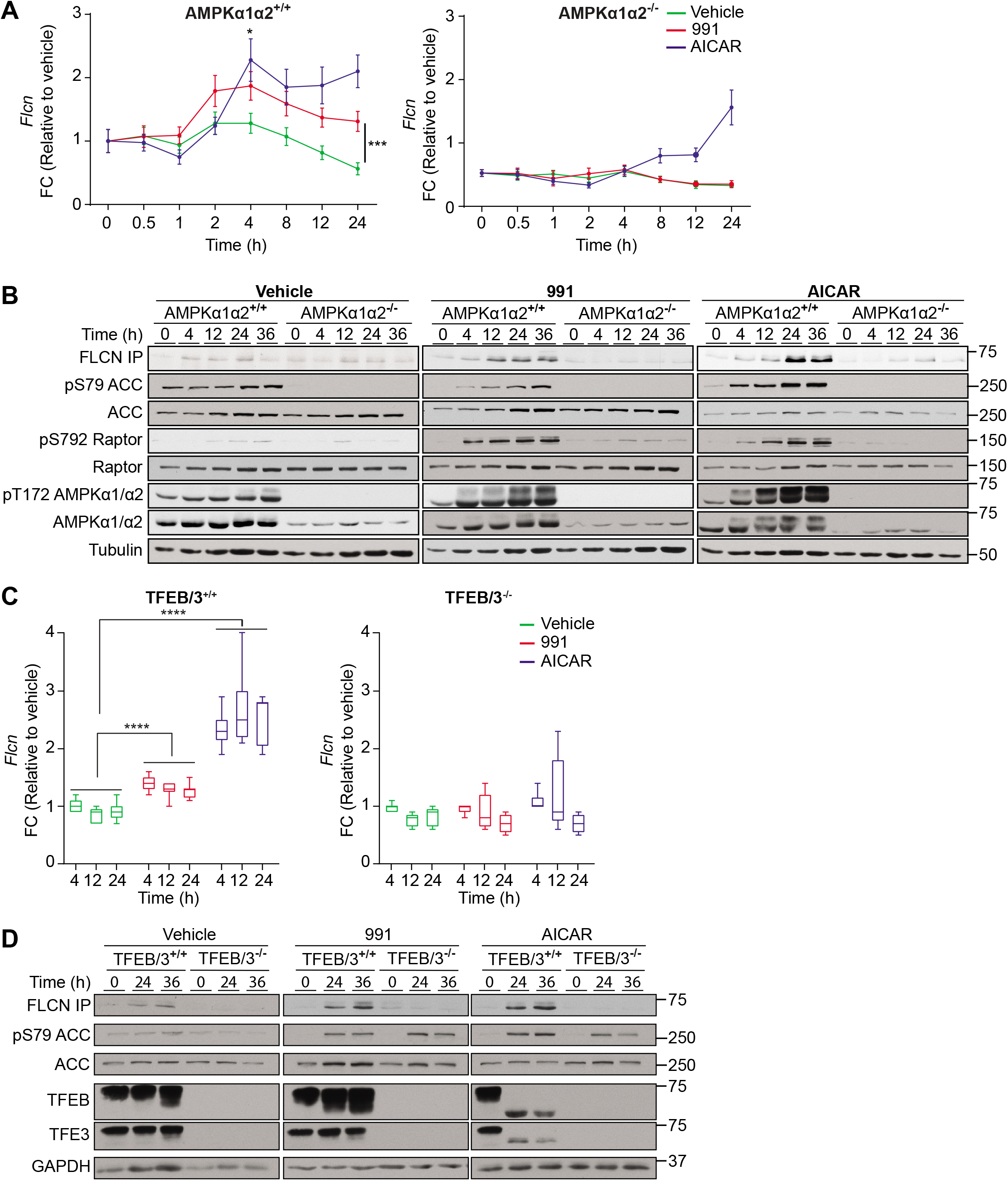
Identification of *Flcn* as an AMPK-regulated gene and TFEB as a transcription factor that mediates this response. (**A**) Relative *Flcn* mRNA quantity was assessed with a Biomark gene expression 192.24 IFC delta gene assay. AMPKα1α2^+/+^ or AMPKα1α2^-/-^ MEF were treated with vehicle (DMSO), 10 μM 991 or 2 mM AICAR, for 0, 0.5, 1, 2, 4, 8, 12 and 24 h. For the analysis, two normalization genes (*Ppia, Tbp*) were used and a two-way ANOVA with interaction was fit to log-transformed data, \**P* < 0.05, \*\*\**P* < 0.001. Each data point represents the mean ± SEM (n=12). (**B**) AMPKα1α2^+/+^ or AMPKα1α2^-/-^ MEF were treated with 10 μM 991 or 2 mM AICAR for the indicated time points. FLCN was immunoprecipitated (IP) from cell lysates and subjected to immunoblot analysis with the indicated antibodies. A representative blot of three independent experiments is shown. (**C**) mRNA level of *Flcn* in TFEB/TFE3 control (TFEB/3^+/+^) or knockout (TFEB/3^-/-^) MEF treated with 10 μM 991 or 2 mM AICAR for 4, 12 and 24 h. Data are presented as a box-and-whisker plots (min to max) of values normalized to control (vehicle-treated cells, 4 h). Data were analyzed by a two-way ANOVA with the factors of time and treatment, plus the interaction between these two factors (n=9). Significance of the treatment factor is indicated, \*\*\*\**P* < 0.0001. (**D**) TFEB/3^+/+^ and TFEB/3^-/-^ MEF were lysed after 0, 24 and 36 h of treatment with 10 μM 991 or 2 mM AICAR. Laemmli extracts (20 μg) were subjected to western blot analysis using the indicated antibodies. FLCN was IP from 500 μg of cell lysate with 1 μg of anti-Folliculin antibody. Images are representative of n=2.

**Table 1.** List of predicted transcription factors mediating AMPK transcriptional response. The 991-responsive genes in mouse embryonic fibroblasts and primary hepatocytes were used to perform an upstream regulator analysis in Ingenuity Pathway Analysis. The table shows the top predicted transcription factors together with their *P-*value.

| Mouse embryonic fibroblasts |  | Mouse primary hepatocytes |  |
| --- | --- | --- | --- |
| Transcriptional regulator | *P*-value | Transcriptional regulator | *P*-value |
| SREBP-1 | 1E-14 | NUPR1 | 2E-08 |
| SREBP-2 | 8E-12 | CREB-1 | 3E-08 |
| STAT4 | 2E-08 | **TFEB** | **6E-08** |
| SIRT2 | 7E-08 | SREBP-1 | 6E-07 |
| SRC2 | 7E-06 | Miz-1 | 1E-05 |
| **TFEB** | **2E-04** | ATF-4 | 1E-04 |
| FOXO3 | 3E-04 | ATF-2 | 3E-04 |
| TP53 | 4E-04 | TCF-3 | 3E-04 |
| NFATc2 | 9E-04 | MITF | 3E-03 |
| IRF-7 | 3E-03 | TAF7L | 1E-02 |
| IRF-3 | 4E-03 | HOXD10 | 1E-01 |
|  |  | HNF-1B | 2E-01 |

### AMPK promotes dephosphorylation and nuclear translocation of TFEB independently of mTOR

It has been reported that the transcriptional activity of TFEB is coupled to its subcellular localization and is regulated by reversible phosphorylation. We thus wanted to determine if AMPK-induced activation of TFEB occurs as a consequence of its dephosphorylation and nuclear localization. To test this hypothesis, we treated WT and AMPK KO MEF with 991 or AICAR for 4 hours and prepared cytoplasmic and nuclear fractions followed by immunoblot analysis (Fig. 6A). ACC was only detected in the cytoplasm as anticipated and its phosphorylation was increased (~2 fold) after treatment with the compounds. In WT MEF both 991 and AICAR similarly decreased and increased TFEB levels in cytoplasmic and nuclear fractions, respectively, whereas the levels of TFEB were not altered in both fractions upon treatment with 991 or AICAR compared to control (vehicle) in AMPK KO MEF. Notably, both compound treatments resulted in a marked increase in a faster-migrating form of TFEB, indicative of TFEB dephosphorylation, in the nuclear fraction in WT but not in AMPK KO MEF (Fig. 6A). To examine if the faster migration of TFEB was associated with dephosphorylation, we generated a phospho-specific antibody against Ser (S)142, one of the key phosphorylation sites impacting cellular localization and activity of TFEB^48^. We verified and confirmed the specificity of the phospho-S142 TFEB antibody by immunoblotting using recombinant WT and non-phosphorylatable Ala (A) mutant form (S142A) of N-terminus Flag-tagged mouse TFEB (Fig. 6B). Given that mTOR has been demonstrated to act as a key upstream regulator of TFEB and that AMPK is known to modulate mTOR activity, we also wanted to address if AMPK-mediated dephosphorylation and nuclear localization of TFEB were dependent on mTOR (Fig. 6C). Consistent with the results shown in Fig. 6A, both 991 and AICAR treatment (for 1 and 4 hours) resulted in a downward band-shift of total TFEB in WT, which was associated with a decrease in S142 phosphorylation. In contrast, compound treatment had no effect on both band-shift and S142 phosphorylation in AMPK KO MEF (Fig. 6C). AICAR induced a robust decrease in mTOR activity as evidenced by a decrease in phosphorylation of p70 S6K (Thr389, a known mTOR-target site) and its downstream substrate ribosomal protein S6 (S6RP) in WT (notably at 4 hours post treatment), but not in AMPK KO MEF. 991 treatment showed only a modest decrease in S6RP phosphorylation in WT, but not in AMPK KO MEF. Treatment with Rapamycin, an mTOR complex 1 (mTORC1) inhibitor, for 1 or 4 hours at three doses (0.05, 0.1, and 0.5 μM) caused a robust decrease in phosphorylation of p70 S6K and S6RP in both WT and AMPK KO MEF. In sharp contrast to this observation, Rapamycin did not lead to any notable band-shift or dephosphorylation (S142) of TFEB in both WT and AMPK KO MEF (Fig. 6C). It has previously been demonstrated that nutrient-induced S142 phosphorylation was resistant to Rapamycin but sensitive to a novel class of mTOR inhibitor (Torin) targeting catalytic site of mTOR^48^. Torin (5 or 10 nM for 1 or 4 hours) inhibited mTOR activity (*i.e.* ablated phosphorylation of p70 S6K and S6RP) to a similar degree compared to Rapamycin. Of note, Torin treatment (10 nM, 4 hours) caused a modest downward band-shift and dephosphorylation (S142) of TFEB, although this was not AMPK-dependent since it was observed in both WT and AMPK KO MEF (Fig. 6D).

**Figure 6.**
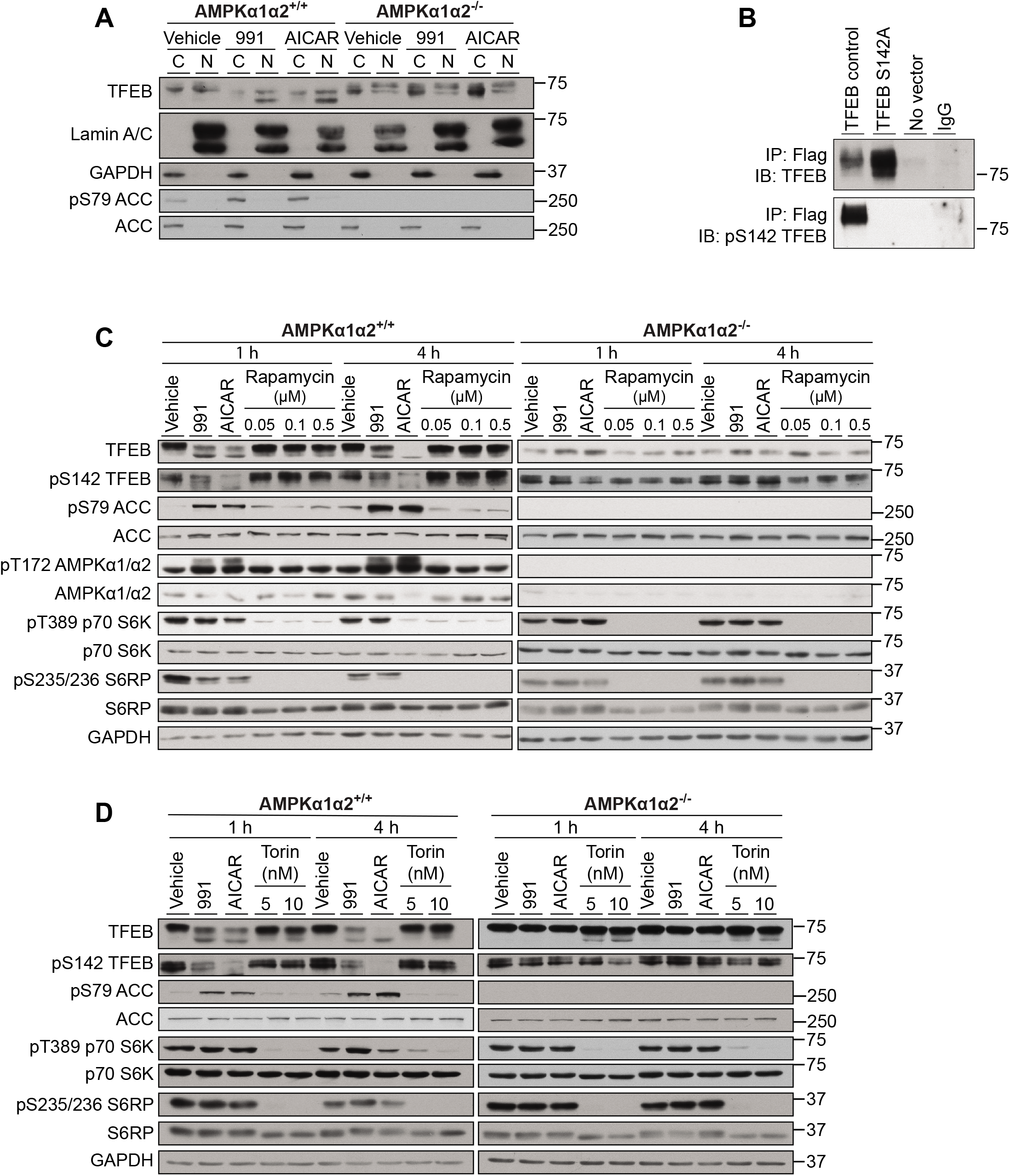
TFEB translocates to the nucleus upon AMPK stimulation independently of mTOR in MEF. (**A**) AMPKα1α2^+/+^ or AMPKα1α2^-/-^ MEF were stimulated with vehicle (DMSO), 10 μM 991 or 2 mM AICAR for 1 h. The cytoplasmic and nuclear fractions were generated using the NE-PER kit accordingly to the manufacturers’ instructions. The lysates (20 μg) were used for western blot analysis of the indicated proteins. (**B**) COS1 cells were transfected for 48 h with no vector (negative control), TFEB control Flag-tagged or Flag-tagged mutated on the phosphorylation site (S142A). TFEB was immunoprecipitated (IP) from 500 μg of cell lysate with 1 μg of anti-Flag antibody, and subjected to western blot analysis (IB) with total TFEB or anti-pS142 TFEB-antibodies. Images are representative of n=2. (**C**) MEF were treated with AMPK activators (10 μM 991 or 2 mM AICAR) or different doses of the mTOR inhibitor Rapamycin (0.05, 0.1 and 0.5 μM) for 1 or 4 h. Cell lysates (20 μg) were subjected to western blot analysis using antibodies against TFEB together with various components of the mTOR- and AMPK-signaling pathways. (**D**) The experiment was performed as described in (**C**), however Torin-2 (5 or 10 nM) was used as an mTOR inhibitor in place of Rapamycin. Figures are representative of n=2.

We next wanted to determine if the AMPK-dependent and mTOR-independent dephosphorylation and nuclear enrichment of TFEB in response to AMPK activators could also be observed in mouse primary hepatocytes (Fig. 7A, B). As anticipated, treatment of hepatocytes with AMPK activators (991, AICAR, and also C13, a recently identified α1-selective AMPK activator^34,49^) for 1 or 4 hours robustly increased phosphorylation of AMPK and ACC in WT, which was totally ablated in AMPK KO hepatocytes (Fig. 7A). AICAR and C13, but not 991, potently reduced phosphorylation of p70 S6K and S6RP in an AMPK-dependent mechanism, except that C13 (4 hours) displayed an AMPK-independent inhibition of mTOR. In control (vehicle-treated) hepatocytes, total TFEB appeared as doublets in WT, whereas in AMPK KO hepatocytes TFEB band was fainter and smeary compared to WT. Consistent with our observations using MEF (Fig. 6A, C and D), AMPK activators resulted in disappearance of the upper band and increased the amount of the lower/faster-migrating form of TFEB in WT, while no apparent change/band-shift in TFEB was observed in AMPK KO hepatocytes. We also observed that treatment of WT hepatocytes with the AMPK activators decreased and increased TFEB levels in cytoplasmic and nuclear fractions, respectively (data not shown). Of note, we attempted to assess TFEB S142 phosphorylation, however it was not detectable most likely due to the low abundance of total TFEB in mouse primary hepatocytes (data not shown). In line with the observations in MEF (Fig. 6C), Rapamycin had no apparent effect on TFEB band-shift/-migration (Fig. 7A). To demonstrate that a loss of band-shift/dephosphorylation of TFEB observed in AMPK KO hepatocytes is intrinsic to AMPK deficiency, we reintroduced AMPK subunits (Flag-α2, β1, γ1) back in AMPK KO cells using adenovirus (Fig. 7B). Genetic deletion of AMPKα1 and α2 catalytic subunit (AMPK KO) resulted in profound reductions in other regulatory subunits (β1 and γ1). When AMPKα2, β1, γ1 were introduced/expressed in the AMPK KO hepatocytes, the effect of 991 (following both 1 and 4 hours) on ACC phosphorylation, as well as a downward band-shift of TFEB, was totally restored. Collectively, we demonstrate that AMPK activators promote dephosphorylation and nuclear localization of TFEB in an AMPK-dependent and mTOR-independent mechanism in MEF and mouse primary hepatocytes.

**Figure 7.**
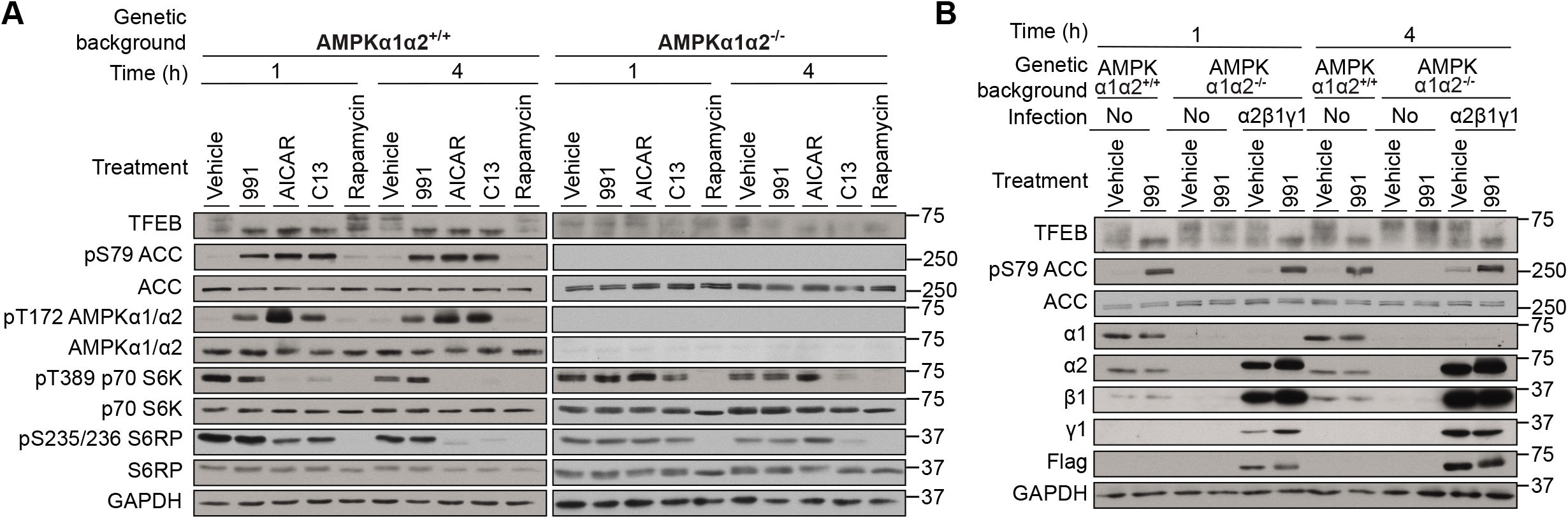
TFEB translocates to the nucleus upon AMPK stimulation independently of mTOR in mouse primary hepatocytes. (**A**) AMPKα1α2^+/+^ or AMPKα1α2^-/-^ mouse primary hepatocytes were isolated and cells were treated with 10 μM 991, 300 μM AICAR, 30 μM C13 or Rapamycin 0.05 μM, for 1 or 4 h. Western blot analysis of the indicated proteins was performed on cell lysates. (**B**) Primary hepatocytes were isolated from AMPKα1α2^-/-^ mice and where indicted were co-infected with three adenoviruses (1:3 MOI) in order to rescue the expression of the three subunits of AMPK (Flag-α2, β1 and γ1). 16 h post infection, the primary hepatocytes were treated for 1 h with vehicle (DMSO) or 10 μM 991. Cell lysates (20 μg) were used for western blot analysis with the indicated antibodies. Figures are representative of n=3.

### *In vivo* evidence that AMPK activation promotes nuclear localization of TFEB and *Flcn* expression using zebrafish

Zebrafish (*Danio rerio*) represents a powerful vertebrate model in biomedical research, given its genetic similarities with humans together with its high fecundity, rapid development and the optical transparency of embryos and larvae. Taking advantage of the zebrafish model, we sought to address if AMPK activation promotes nuclear localization of TFEB at organismal/tissue levels and across species. To this end, we employed zebrafish transgenically-expressing zebrafish Tfeb fused to ZsGreen (ZsGreen-Tfeb), as well as mCherry fused to a nuclear localization signal (NLS) under the actin alpha cardiac muscle 1b *actc1b* promoter, which drives gene expression in skeletal muscle. Fluorescence-based confocal microscopy was performed on embryos 3 days post fertilization following treatment with vehicle (DMSO) or 991 (10 μM) for 24 hours (Fig. 8A). We verified that 991 (10 μM for 24 hours) increases the phosphorylation of AMPK and ACC by immunoblot analysis using larvae protein extracts (data not shown). Under vehicle-treated condition, the ZsGreen-Tfeb was distributed throughout the entire cell likely due to cytosolic localization of Tfeb. In support of this, merging the ZsGreen-Tfeb with the mCherry-NLS images showed that there was no apparent nuclear localization of Tfeb (Fig. 8A). We observed a change in the distribution pattern of ZsGreen-Tfeb going from a diffuse pattern to a forming puncta within the cell. Merging this image with the mCherry-NLS shows that these puncta superimpose with the signal from the mCherry-NLS image. This suggests that treatment of zebrafish larvae with 991 leads to translocation of ZsGreen-Tfeb to the nucleus, which is consistent with results obtained with cell fractionation and immunoblot analysis in MEF (Fig. 6A). Altogether, these results compellingly demonstrate AMPK plays an important role in the regulation and activation of TFEB across species.

**Figure 8.**
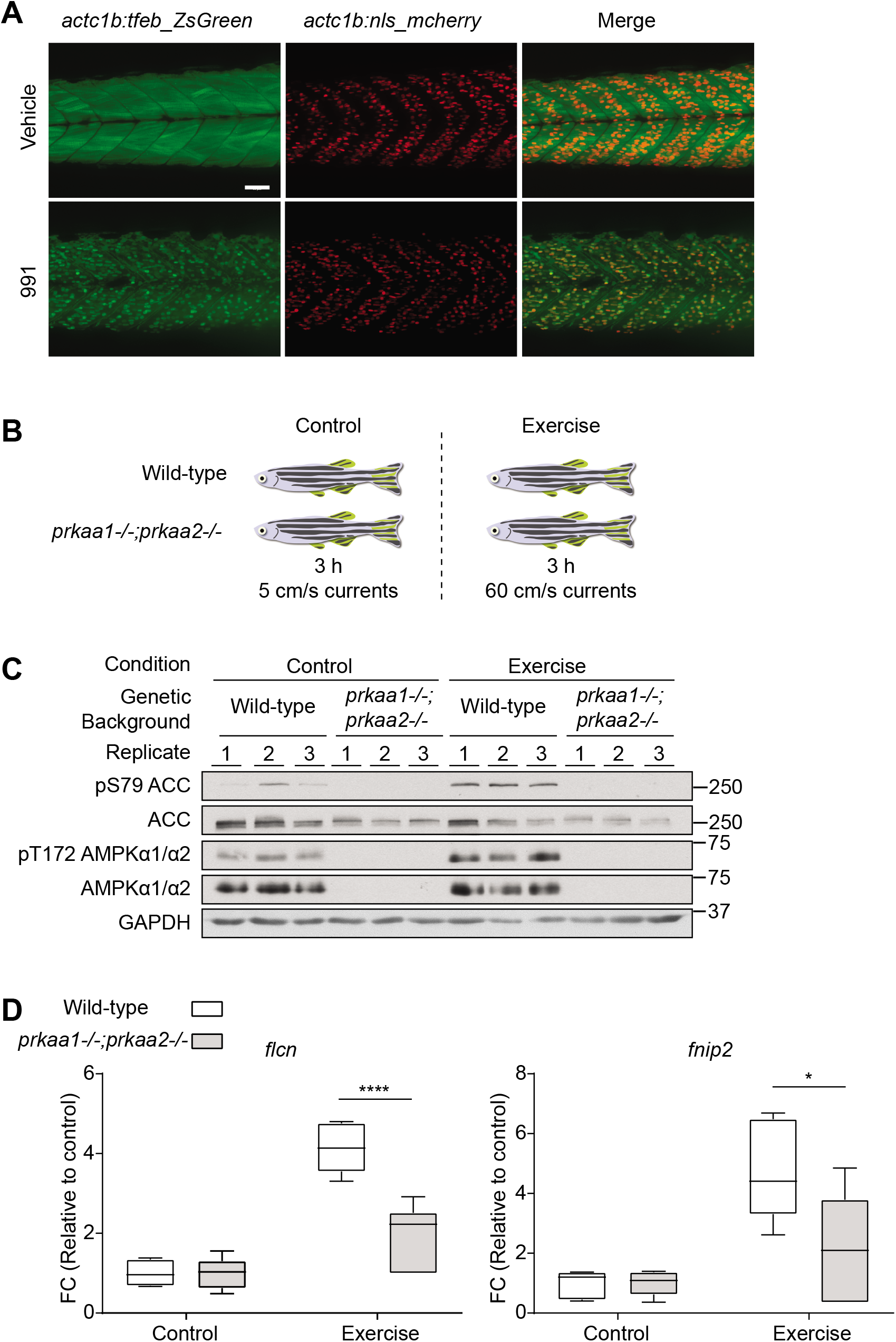
Activation of AMPK in zebrafish leads to translocation of TFEB to the nucleus and increased expression of *flcn* and *fnip2*. (**A**) *Tg(actc1b:tfeb-ZsGreen);Tg(actc1b:nlsmCherry)* embryos at 3 dpf were treated with 991 10 μM or vehicle (DMSO) for 24 h. Embryos were mounted live in water containing 0.016% tricaine and imaged with ImageXpress confocal system at 20X magnification, the scale bar corresponds to 50 μm. (**B**) The figure summarizes the experimental scheme implemented to acutely exercise wild-type and *prkaa1*^-/-^*;prkaa2*^-/-^ zebrafish. An electronic controller and a motor-driven propeller were used to adjust the water velocity to maintain the indicated speed. (**C**) Immediately after the three hours of training, muscle samples were collected from the zebrafish. Tissue lysates (20 μg) were separated by SDS-PAGE and western blot analysis was performed using the indicated antibodies. (**D**) mRNA transcript levels of *flcn* and *fnip2* were calculated using *ef1a* and *nrf1* as reference genes. Data are shown as box-and-whisker plots (min to max) of values normalized to control wild-type (n=6). Two-way ANOVA with the factors of genetic background and exercise, plus the interaction between these two factors was performed. The graph shows the significance of the interaction, \**P* < 0.05, \*\*\*\**P* < 0.0001.

Having demonstrated that pharmacological activation of AMPK promotes nuclear localization of TFEB and its transcriptional activation leading to induced *Flcn* expression, we wanted to study if a physiological activation of AMPK, such as physical exercise, would also increase *Flcn* expression *in vivo*. For this purpose, we generated loss of function of *prkaa1/prkaa2* (which encodes paralogue of human/mouse AMPKα1 and α2, respectively) double KO zebrafish using CRISPR-Cas9 genome editing (Parisi *et al.*, in preparation). Using this model, we investigated whether expression of *Flcn* and *Fnip2*, the paralogue of *Fnip1*, alters in response to an acute bout of swimming exercise. We initially established an acute exercise training protocol which stimulates AMPK in zebrafish (as summarized in Fig. 8B). The control and exercise groups comprised a mixed population of WT and knockout fish, to ensure consistency between the experimental groups. The control group was placed in the swim tunnel and subjected to a low flow speed (5 cm/s) for 3 hours. For the exercise group, an acclimatization period of 20 min at a low flow speed (5 cm/s) was followed by a gradual increase of water velocity up to 80% of maximum speed capacity (*i.e.* 60 cm/s for this cohort of fish), which was maintained for approximately 2 hours. Notably, the maximum speed of the fish, defined as the maximum sustainable swimming speed, was determined in a previous grouped endurance test following the existing guidelines^50^. Subsequently, the skeletal muscles were rapidly extracted to avoid AMPK activation due to cellular stress (*e.g.* hypoxia). The loss of expression/function of AMPK in the knockout fish was assessed by immunoblotting, which confirmed the absence of expression and activity of the AMPK catalytic subunits (Fig. 8C). We confirmed that an acute bout of exercise increased AMPK activity as judged by an increase in phosphorylation of AMPK and ACC in WT muscle (Fig. 8C). Finally, we compared by qPCR the levels of *flcn* and *fnip2*, the paralogue of *fnip1*. The results showed an increased level of expression of both genes after physical exercise in WT zebrafish, and that this increase was significantly decreased in *prkaa1/prkaa2*-deficient zebrafish (Fig. 8D). Taken together, these data suggest that exercise increases the activity of AMPK in zebrafish, which is associated with an increase in *flcn* and *fnip2* expression that is at least in part mediated through AMPK.

## DISCUSSION

It is well documented that AMPK not only elicits a plethora of acute metabolic responses, but also promotes metabolic reprogramming by modulating gene expression through regulation of specific transcription factors and transcriptional co-activators^12,27,51^. In the current study, we performed whole-genome transcriptome profiling in AMPK-intact or -deficient mouse embryonic fibroblasts and hepatocytes. We identified several new AMPK-dependent/-regulated genes and pathways that are differentially regulated in a cell type- and a compound-specific manner. The major finding of this study was that we found TFEB, a transcription factor and emerging regulator of lysosomal biogenesis/autophagy^52^, as one of the key mediators of AMPK-dependent transcriptional responses.

We performed transcriptome profiling using two different AMPK activators (AICAR and 991), known to target distinct regulatory sites/mechanisms. AICAR is a valuable and the most commonly used pharmacological AMPK activator, which significantly contributed to uncover critical metabolic functions of AMPK for decades^53–55^. In recent years, with the development and availability of genetic technologies/tools (*i.e.* AMPK KO models), as well as highly-specific AMPK activators targeting ADaM site, the specificity of AICAR has been questioned. Several studies have reported AICAR’s off-target effects^17^, for example on AMP-regulated enzymes such as fructose 1,6-bisphosphatase, a key regulator for hepatic gluconeogenesis^56,57^. To our knowledge, this is the first study investigating the effect of AICAR on genome-wide gene expression employing AMPK-deficient cell models to dissect AMPK-dependent/-independent effects. It was striking to find out that the vast majority of the transcripts were significantly altered with AICAR in the absence of functional AMPK. This implied that previously claimed signaling, cellular, as well as physiological effects induced by AICAR may not be mediated through AMPK (unless its off-target effects were ruled out using AMPK KO models as a control). In contrast, 991 exhibited nearly exclusive specificity for targeting AMPK in both MEF and hepatocytes consistent with the specificity of this compound that we showed *in vitro* (cell-free) in our previous study^4^. 991 activates both β1- and β2-containing complexes (with higher affinity to the β1 complexes), thus it acts as a pan/total AMPK activator. Given that isoform-specific AMPK activators have recently been identified^34,58^, it would be of interest to elucidate isoform-specific gene responses and metabolic programming.

The cellular localization and activity of TFEB are primarily regulated by its phosphorylation status. Two serine residues, S142 and S211, in the TFEB have mainly been proposed to play a key role in determining its subcellular localization^48,52^. When both sites are phosphorylated, TFEB is kept inactive in the cytosol, while mutation of either S142 or S211 to non-phosphorylatable alanine residue rendered TFEB constitutively active through keeping it within the nucleus. mTOR and ERK2 are the main protein kinases known to phosphorylate TFEB under nutrient-rich conditions supplemented cells with conventional cell culture medium) in most cell types. AMPK inhibits mTOR through multiple mechanisms^59–61^, and it has recently been reported that AMPK controls endolysosomal function through suppression of mTOR and its subsequent regulation of TFEB^42^. Therefore, we initially hypothesized that 991- or AICAR-induced activation of TFEB, via dephosphorylation and nuclear localization, is mediated through the AMPK-dependent inhibition of mTOR. Contrary to our hypothesis, we observed that acute treatment with a specific AMPK activator, 991^4,20^, promoted TFEB dephosphorylation in the absence of detectable inhibition of mTOR (as judged by phosphorylation of p70 S6K and S6RP) in primary hepatocytes. In a previous study, it has been shown that phosphorylation of TFEB at S142 represents a Rapamycin-resistant, but Torin-sensitive site^48^. In line with this, we observed that the S142 phosphorylation was not affected by Rapamycin. However, in contrast to the previous observation, Torin displayed only a marginal effect on TFEB dephosphorylation (*i.e.* appearance of a minor faster-migrating form of the total TFEB and reduced S142 phosphorylation), and notably this was not only observed in WT, but also in AMPK KO MEF. Collectively, in MEF and primary hepatocytes, 1) we have demonstrated that acute inhibition of mTOR (*i.e.* 1-4 hours) does not modulate TFEB phosphorylation, and 2) 991-induced dephosphorylation of TFEB is unlikely to be mediated through AMPK-dependent suppression of mTOR. However, it would be important to assess if other phosphorylation sites (*e.g.* S211, S138), which are proposed to play key roles for cellular distribution of TFEB^62^, are regulated upon AMPK activation.

The mechanism by which AMPK dephosphorylates and activates TFEB is unknown. It has been demonstrated that nutrient deprivation induces the release of lysosomal Ca^2+^ through Ca^2+^ channel mucolipin 1. This activates calcium/calmodulin-activated serine/threonine phosphatase calcineurin (also known as protein phosphatase 2B), which binds to and dephosphorylates TFEB, thus promoting its nuclear localization and autophagy induction^63^. It would be interesting to determine if AMPK-mediated activation of TFEB regulates the catalytic activity of calcineurin and/or interaction between TFEB and calcineurin. It has also been shown that nutrient/glucose deprivation-induced AMPK activation regulates lysosomal and autophagy gene expression through phosphorylation at S659 and nuclear localization of acetyl-CoA synthetase 2 (ACSS2)^64^. Phosphorylated ACSS2 forms a complex with TFEB, which modulates lysosomal and autophagosomal genes by locally producing acetyl CoA for histone H3 acetylation in the promoter regions of these genes. Therefore, AMPK indirectly modulates the transcriptional activity of TFEB after its nuclear translocation via inhibition of mTOR in response to energy stress. Nonetheless, in the current study we found *Acss2* as one of the AMPK-dependent genes upregulated at mRNA levels in response to both 991 and AICAR. Whether this upregulation of *Acss2* is linked to an increase in its protein/phosphorylation levels needs to be determined.

Although TFEB is an established master regulator of lysosomal biogenesis^65,66^, emerging evidence suggests that it also acts as key controller for various other cellular and metabolic responses, including lipid metabolism in liver^67^, mitochondrial biogenesis in muscle^44^, as well as modulation of the immune response^68^. In support of this, we identified genes that are involved in lipid/cholesterol signaling and metabolism (*Acss2, Crebrf, Hmgcr, Ldlr, Lpin1, Msmo1, Pde4b*) and immunity (*Ifit1, Tollip*). It would be of interest to determine if these genes are regulated through the AMPK-TFEB axis, as we have shown for *Flcn*. The tumor suppressor FLCN, responsible for the Birt-Hogg Dubé renal neoplasia syndrome (BHD), is an AMPK-interacting partner which has recently been proposed to function as a negative regulator of AMPK^46,47^. It has been reported that ablation of FLCN expression or loss of FLCN binding to AMPK cause constitutive activation of AMPK, which was associated with enhanced osmotic stress resistance and metabolic transformation. Notably, genetic inactivation of FLCN in adipose tissue led to a metabolic reprogramming characterized by enhanced mitochondrial biogenesis and browning of white adipose tissue^45^. Mechanistically, adipose-specific deletion of FLCN results in induction of the PGC-1 transcriptional coactivator through relieving mTOR-dependent cytoplasmic retention of TFE3^69^ and/or activation of AMPK^45^. It has been shown that exercise promotes TFEB translocation into the myonuclei, which regulates glucose/glycogen metabolism by controlling expression of glucose transporters, glycolytic enzymes, as well as pathways linked to glucose homeostasis^44^. Moreover, muscle-specific over-expression of TFEB mimics the effects of exercise training and promotes metabolic reprogramming through induction of gene expression involved in mitochondrial biogenesis and function. We showed *in vivo* using zebrafish that exercise induces *Flcn* and *Fnip2* expression at least partially through an AMPK-dependent mechanism. Moreover, we proved that activation of AMPK by 991 induces Tfeb translocation to the nucleus, suggesting that the increased gene transcription of *Flcn* and *Fnip2* observed after exercise are likely mediated through the ability of AMPK to promote Tfeb translocation in skeletal muscle. Whether FLCN mediates part of metabolic responses downstream of TFEB/TFE3 or increased expression of FLCN functions as negative feedback loop to suppress AMPK to avoid its prolonged activation of AMPK is unknown.

Coincidentally, we noticed that in the dataset generated by Cokorinos *et al*, the mRNA levels of *Flcn* and *Fnip1/2* were elevated in response to acute or chronic treatment with the ADaM site-binding allosteric AMPK activator PF-739 in mouse skeletal muscle^25^. This strongly supports the findings from our current study that activation of AMPK leads to changes in *Flcn* and *Fnip1*/*2 in vivo* across multiple species and tissues. Our study suggests that the ability of PF-739 to increase transcription of *Flcn* and *Fnip1/2* are likely mediated by its ability to regulate TFEB.

In summary, we demonstrated in fibroblasts, hepatocytes, as well as at whole organism levels, *in vivo* using zebrafish, that pharmacological/physiological activation of AMPK promoted nuclear translocation of TFEB. This appeared to be through an apparent effect on dephosphorylation of TFEB, independent of mTOR, and was associated with induction of a tumor suppressor FLCN through activation of its promoter activity. Future studies using gain of function models of FLCN in skeletal muscle and other tissues (*e.g.* liver) could reveal the physiological significance of the AMPK-TFEB-FLCN pathway.

## MATERIALS AND METHODS

### Materials

The materials used comprise 5-aminoimidazole-4-carboxamide riboside (AICAR, OR1170T, Apollo Scientific), Protein G Sepharose (P3296; Sigma), 991 (5-[[6-chloro-5-(1-methylindol-5-yl)-1H-benzimidazol-2-yl]oxy]-2-methyl-benzoic acid, CAS number 129739-36-2)^31^, Rapamycin (R0395, Sigma) and Torin-2 (SML1224, Sigma). General and specific cell culture reagents were obtained from Life Technologies. All other materials unless otherwise indicated were from Sigma.

### Antibodies

Total FLCN antibody was purchased from Proteintech (#11236-2-AP). Flag (#F7425), α-tubulin (#T6074), and GAPDH (#G8795) were obtained from Sigma. AMPKα1 (#07-350) and AMPKα2 (#07-363) antibodies were obtained from Merck Millipore. Acetyl-CoA carboxylase (ACC; #3676), phospho-ACC1 (Ser79; #3661), AMPKα (#2532), phospho-AMPKα (Thr172; #2535), AMPKβ1 (#4178), lamin A/C (#4777), p70 S6 kinase (#9202), phospho-p70 S6 kinase (Thr389; # 9206), raptor (#2280), phospho-raptor (Ser792; #2083), S6 ribosomal protein (#5G10), phospho-S6 ribosomal protein (Ser235/236; #2211) and TFE3 (#14779) antibodies were obtained from Cell Signaling Technology. TFEB (#A303-673A) antibody purchased was from Bethyl Laboratories. Horseradish peroxidase-conjugated secondary antibodies were from Jackson ImmunoResearch Europe. AMPKγ1 (#TA300519) antibody was from OriGene. Site-specific rabbit polyclonal antibody against phospho-TFEB (Ser142) was generated by YenZym Antibodies (South San Francisco, CA, USA) by immunisation with a phosphorylated peptide of the sequence identical between human and mouse was used (*i.e.* PN-*S-PMAMLHIGSNPC-amide, where the prefix *denotes the phosphorylated residue).

### Cell culture

Mouse embryonic fibroblasts (MEF) from wild-type (WT) and AMPK α1^-/-^/α2^-/-^ mice were generated as described previously^70^. TFEB^-/-^/TFE3^-/-^ MEF were a kind gift from Rosa Puertollano (NIH). MEF were cultured in DMEM-Glutamax supplemented with 10% fetal calf serum (FCS) and 1% penicillin streptomycin. Cells were seeded at ~80% confluency and treated the following day at the indicated treatments described in the figures. Cells were washed with ice-cold PBS and scraped into lysis buffer (50 mM HEPES, 150 mM NaCl, 100 mM NaF, 10 mM Na-pyrophosphate, 5 mM EDTA, 250 mM sucrose, 1 mM DTT, 1% Triton X-100, 1 mM Na-orthovanadate, 0.5 mM PMSF, 1 mM benzamidine HCl, 1 μg/ml leupeptin, 1 μg/ml pepstatin-A, 1 mM microcystin-LR). Primary hepatocytes were isolated AMPKα1/α2 liver-specific knockout mice and control AMPKα1^lox/lox^α2^lox/lox^ mice littermates (10-week-old males) by collagenase perfusion and cultured as previously described ^35^. The experiments were performed accordingly with the European guidelines (approved by the French authorisation to experiment on vertebrates (no. 75-886) and the ethics committee from University Paris Descartes (no. CEEA34.BV.157.12). Briefly the cells were plated in M199 medium containing Glutamax and supplemented with 100 U/ml penicillin, 100 μg/ml streptomycin, 10% (v/v) FBS, 500 nM dexamethasone (Sigma), 100 nM triiodothyronine (Sigma), and 10 nM insulin (Sigma). The hepatocytes were allowed to attach (4 hours), and were then maintained in M199 medium with antibiotics and 100 nM dexamethasone for 16 hours. Experiments were performed the following morning by treating hepatocytes with the indicated compounds (*e.g.* AMPK activators, mTOR inhibitors) for 1 or 4 hours. For viral infection, the hepatocytes were co-infected (1:3 MOI) with three individual adenovirus encoding AMPK subunits α2, β1, γ1 into AMPK α1^-/-^/α2^-/-^ hepatocytes to restore expression of the AMPK trimeric complexes. 16 hours following adenovirus infection, the cells were treated for 1 h with vehicle (DMSO) or 991 (10 μM). Following the above described treatments, media were aspirated and cells lysed on ice in cold lysis buffer. Lysates were snap-frozen in liquid nitrogen and stored at −80°C for subsequent analyses. Lysates were clarified at 3500 g for 15 min at 4 °C and quantified using Bradford reagent and BSA as standard.

### RNA extraction, microarray and bioinformatic analysis

Total RNA was extracted from cells using RNAdvance Tissue Kit (A32645; Beckman Coulter) and quantified by RiboGreen (R11490; ThermoFisher). RNA integrity was determined by Fragment Analyzer, (DNF-471, Advanced Analytical) and an RNA Quality Number greater than 8 was observed. 300 ng of total RNA were used as input to produce labeled cRNA targets with the TotalPrep-96 RNA amplification kit, following manufacturer’s instructions. cRNA quality was analyzed by Fragment Analyzer, subsequently 11 μg of cRNA was fragmented and hybridized onto Affymetrix Gene Chip Mouse 430 2.0 following manufacturer’s instructions. Partek Genomics Suite software was used to analyze the data from CEL files. Values were normalized using Robust Multichip Average (RMA) method. The removal batch effect was applied during the analysis of the hepatocytes samples. Based on the normal distribution of the datasets, the parametric Pearson’s product moment correlation was applied for quality control. Two-way analysis of variance (ANOVA) with Benjamini & Hochberg multiple testing correction was applied to discriminate 991 versus control and AICAR versus control conditions. The moderated *P*-value was set at 0.05 for the interaction within genetic background and treatment as well as for the pairwise comparisons. In addition, a fold-change cutoff of 1.3 was applied. Considering the quality test results, two samples emerged as outlier and therefore were excluded from the analysis (*i.e.* MEF WT, 991-treated, technical replicate 1 and hepatocytes AMPK α1^-/-^/α2^-/-^, vehicle-stimulated, technical replicate 4, biological replicate 1). The gene ontology was performed using DAVID ^37,38^ and the transcription factor prediction using the upstream regulator analysis available in Ingenuity pathway analysis^39^ (QIAGEN Inc., https://www.qiagenbioinformatics.com/products/ingenuity pathway-analysis).

### RT-qPCR

For cDNA synthesis, 500 ng of total RNA was used as starting material for the PrimeScript RT Kit (Takara, #RR037A). RT-qPCR reactions were performed on a LightCycler 480 with SYBR Green Assay (Roche, #04707516001), with primers at a final concentration of 0.3 μM/reaction. All the primers used in this study are summarized with sequences in Supplementary Table 2. For mouse samples, three normalization genes were used, namely Acyl-CoA synthetase short-chain (*Acss*), β2 microglobulin (*B2m*), Peptidylprolyl isomerase-a (*Ppia*), and their stability was assessed using GeNorm (M value < 0.6)^71,72^. For zebrafish samples, two normalization genes were used, being elongation factor 1-α (*ef1a*) and nuclear respiratory factor 1 (*nrf1*). Normalized values were calculated by dividing the average expression value by a factor equal to the geometric mean of the normalization genes^71^.

The relative *Flcn* mRNA quantity of the samples used in Fig. 3D was assessed using a Biomark gene expression 192.24 IFC delta gene assay, Fluidigm Biomark, following manufacturer’s instruction. Ct values were calculated using the system’s software (Biomark Real-time PCR analysis, Fluidigm). For the analysis, two normalization genes were used (*Ppia* and TATA box-binding protein (*Tbp*)), afterwards two-way ANOVA was fit to the log-transformed data.

### Generation of *actc1b:tfeb-ZsGreen; actc1b:nls-mCherry* double transgenic fish

Adult AB zebrafish were raised at 28°C under standard husbandry conditions. All experimental procedures were carried out according to the Swiss and EU ethical guidelines and were approved by the animal experimentation ethical committee of Canton of Vaud (permit VD3177). Transgenic zebrafish *Tg(actc1b:tfeb-ZsGreen)*^*nei08*^ and *Tg(actc1b:nls-mCherry)*^*nei09*^ were independently generated using I-SCEI meganuclease mediated transgenic insertion into one-cell stage embryos as previously described^73^. One founder for each transgenic line was selected and subsequent generations were propagated and expanded. The two lines were crossed to generate double transgenic embryos, which have been raised at 28°C under standard laboratory conditions before treatment. Double transgenic embryos were selected at 3 dpf and treated with 991 10 μM or DMSO in 96 well plates (n=12). After 24 h of treatment embryos were anesthetized with 0.016% tricaine and imaged with ImageXpress confocal system at 20X magnification (Molecular Devices). Z stack images were captured for each embryo and maximal projection images were produced.

### Generation of *prkaa1*^-/-^*;prkaa2*^-/-^ double knockout fish

The *prkaa1*^-/-^*;prkaa2*^-/-^ double knockout fish were generated by using the CRISPR/Cas9 approach (manuscript in preparation). Two gRNA targeting *prkaa1* exon 5 and *prkaa2* exon 6 were designed using the Chopchop online tool (http://chopchop.cbu.uib.no/index.php) and ordered as DNA gene strings. After PCR amplification and purification, they were used as input for in vitro transcription with the MEGAshortscript T7 Kit (Thermo Fisher Scientific). Subsequently the transcripts were purification with the RNA Clean & Concentrator kit (Zymo Research) and their concentration was determined by Nanodrop. *prkaa1*^-/-^ and *prkaa2*^-/-^ single mutants were generated by independent co-injection of single cell stage AB embryos with 50 pg of gRNA and 200 pg of the GeneArt™ Platinum™ Cas9 Nuclease (Thermo Fisher Scientific). Injected embryos were grown to adulthood and outcrossed with AB wild-type fish to identify individuals with mutant germline. Putative filial 1 mutants were fin clipped, their genotype was determined by HRM qPCR (see Supplementary Table X for primers sequence) and the mutation identified by sequencing. For *prkaa1*, we selected a 5 bp deletion and for *prkaa2* a 7 bp deletion, both resulting in frameshift and premature stop codons. Single heterozygous animals for the same gene were in-crossed to obtain single *prkaa1*^-/-^ or *prkaa2*^-/-^. The homozygotes were then crossed to obtain double heterozygous fish (*prkaa1*^+/-^;*prkaa2*^+/-^). Finally, double knockout animals *prkaa1*^-/-^;*prkaa2*^-/-^ were obtained by double heterozygous incross and their genotype was determined by HRM qPCR.

### Acute exercise protocol

Acute exercise was performed using a 5 L swim tunnel by Loligo^®^ Systems (#SW10050). The day before the acute exercise session, 6 wild-type and 6 *prkaa1*^-/-^;*prkaa2*^-/-^ 4 months old siblings were subjected to an endurance test to determine their exercise speed. Baseline critical speed (Ucrit) was determined as the speed by which the weakest zebrafish fatigued (5 sec at the rear of the tunnel). Based on this result, 6 wild-type and 6 *prkaa1*^-/-^;*prkaa2*^-/-^ fish were let habituate for 20 min at a low current speed of 10 cm/sec and then trained for 2 h 40 min at 55 cm/sec (75% of baseline Ucrit). As control group, the 6 wild-type and 6 *prkaa1*^-/-^;*prkaa2*^-/-^ fish were placed in the swim tunnel for 3 h at a low swim speed of 10 cm/s. At the end of the 3 h, fish were euthanized immediately and trunk muscle was isolated, flash frozen in liquid nitrogen and processed for RNA and protein extraction.

### Bicistronic luciferase assay

MEF were transfected with Lipofectamine 3000 (L3000001, ThermoFisher), 12 h post transfection cells were treated with 30 μM 991 for 12 h. After the treatment, cells were washed once with PBS and harvested in Glo Lysis Buffer (Promega, #E2661). Cell lysates were centrifuged at 3500 rpm for 10 min at room temperature. Luciferase assay was measured using a Dual-Luciferase Reporter Assay System STOP and GLO kit (Promega, #E1910). Differences in the ratio of Firefly to NanoLuc luciferase signals were analyzed for statistical significance by 2-way ANOVA.

### Immunoprecipitation and immunoblotting

For immunoprecipitation of FLCN, 200 μg of lysates were incubated with 1 μg of antibody and 5 μl of Protein-G Sepharose, on a shaker (1000 rpm), overnight at 4°C. Immunoprecipitates or total lysates were first denatured in SDS sample buffer, separated by SDS-PAGE, and then transferred to nitrocellulose membrane. Membranes were blocked for 1 h in 10 mM Tris (pH 7.6), 137 mM NaCl, and 0.1% (v/v) Tween-20 (TBST) containing skimmed milk 5% (w/v). Membranes were incubated in primary antibody prepared in TBST containing 1% (w/v) BSA overnight at 4°C.

### Cloning and mutagenesis

All plasmid constructs were generated using standard molecular biology techniques. The promoter sequences and the TFEB/3 binding site were identified from the EPD promoter database (https://epd.vital-it.ch/index.php). The promoter regions were amplified from mouse genomic DNA (Promega), then ligated into a modified pNL 1.2 luciferase vector (Promega). The luciferase assay was carried out according to the company’s protocol (Promega).

To express transiently the mouse TFEB cDNA, the clone MR223016 was ordered from Origene. The phospho sites corresponding to human S122, S142 and S211 were mutated to alanine using standard molecular biology methods. The sequences of all constructs were verified in house utilizing the BigDyeR Terminator 3.1 kit and the 3500XL Genetic analyzer (ABI-Invitrogen).

### Statistical analysis

For the statistical analysis, two-way ANOVA with interaction was used to analyze the data or was fit to the log-transformed data, as specified in the figure legend. Data are expressed as boxand-whisker plot (min to max), log2 fold-change of the mean ± SEM or SD, as indicated. Differences between groups were considered statistically significant when *P*-value < 0.05.

### Accession numbers

The microarray data were deposited in GEO, the accession number will be publicly available once the article is accepted.

## ACKNOWLEDGEMENTS

We thank Dr Rosa Puertollano (National Institute of Health, Bethesda, MD) for her generous gift of the TFEB^-/-^/TFE3^-/-^ mouse embryonic fibroblasts. We thank Dr Julien Marquis for his technical advice on qPCR execution and analysis and Dr Giulia Lizzo for her advice on the gene ontology analysis. We also thank Dr Matthew Sanders for his critical review and editing of the manuscript.

## AUTHOR CONTRIBUTIONS

K.S., P.D. and C.C. designed the study. C.C. performed all the treatments in MEF and the promoter activity assay. C.C. also prepared and analyzed MEF, zebrafish and hepatocytes samples, generated the figures, contributed to analysis and data interpretation. M.F. performed the hepatocytes isolation, viral infection, compound treatments, cell collecting and subcellular fractionation. M.D. performed molecular cloning and mutagenesis on TFEB and FLCN promoter. L.B. supervised C.C. during the setup of the gene expression study. S.M. executed the microarray and contributed to mRNA preparation. B.V. and M.F. generated liver-specific AMPK KO mouse model and AMPK KO MEF. B.V. assisted the design of the hepatocytes study. G.L. contributed to the statistical analysis performing ANOVA analysis fit to log-transformed data. F.R. analyzed with C.C. the microarray data. A.P. generated the AMPK *prkaa1*^-/-^*;prkaa2*^-/-^ zebrafish line and performed with C.C. the fish exercise. G.C. generated and performed experiments on the *actc1b:tfeb-ZsGreen;actc1b:nls-mCherry* double transgenic zebrafish. P.G. supervised A.P. and G.C. and contributed study design and interpretation of zebrafish studies. P.D. supervised the microarray and qPCR experiments design, execution and analysis. K.S and P.D. supervised C.C. C.C. and K.S. wrote the manuscript and all the authors contributed to writing/editing the manuscript.

## CONFLICT OF INTEREST

G. Civiletto, C. Collodet, M. Deak, P. Descombes, P. Gut, G. Lefebvre, S. Metairon, A. Parisi, F. Raymond and K. Sakamoto are full time employees of Nestlé Research (Switzerland).

---

**Supplementary Figure 1.**
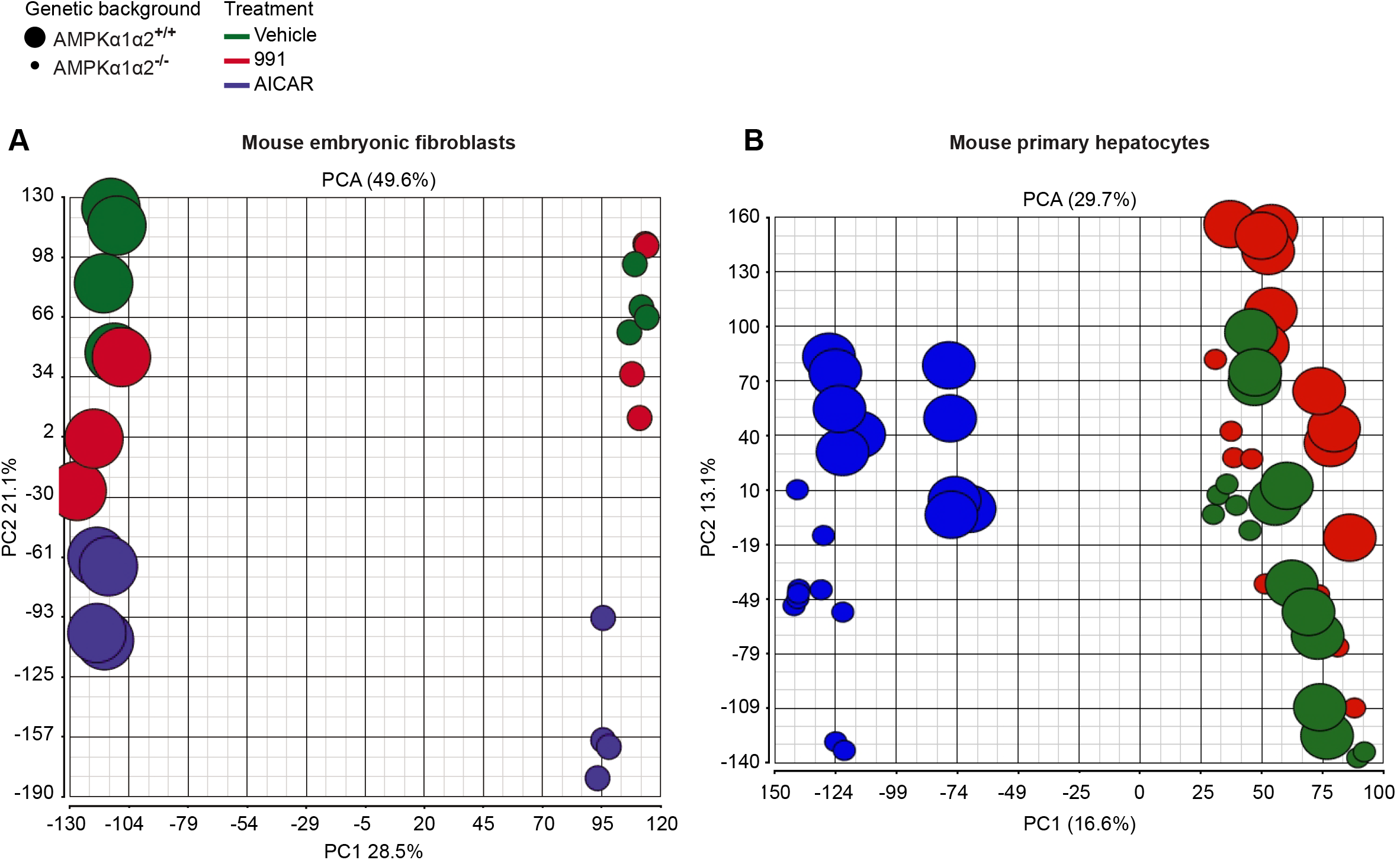
Principal component analysis (PCA) score plots. (**A**) PCA score plot based on gene expression dataset generated from mouse embryonic fibroblasts treated with vehicle (DMSO), 10 μM 991 or 2 mM AICAR for 4 h. The first two principal components (PC1 and PC2) explain 28.5% and 21.1% of total variance in the data, due to genetic background and treatment, respectively. (**B**) PCA score plot obtained from the gene expression dataset of mouse primary hepatocytes treated with vehicle (DMSO), 3 μM 991 or 300 μM AICAR for 4 h. AICAR and 991 treatment are responsible for 16.6% and 13.1% of the total variance in the data, represented by the two principal components (PC1 and PC2). Samples are depicted as indicated in the legend.

**Supplementary Figure 2.**
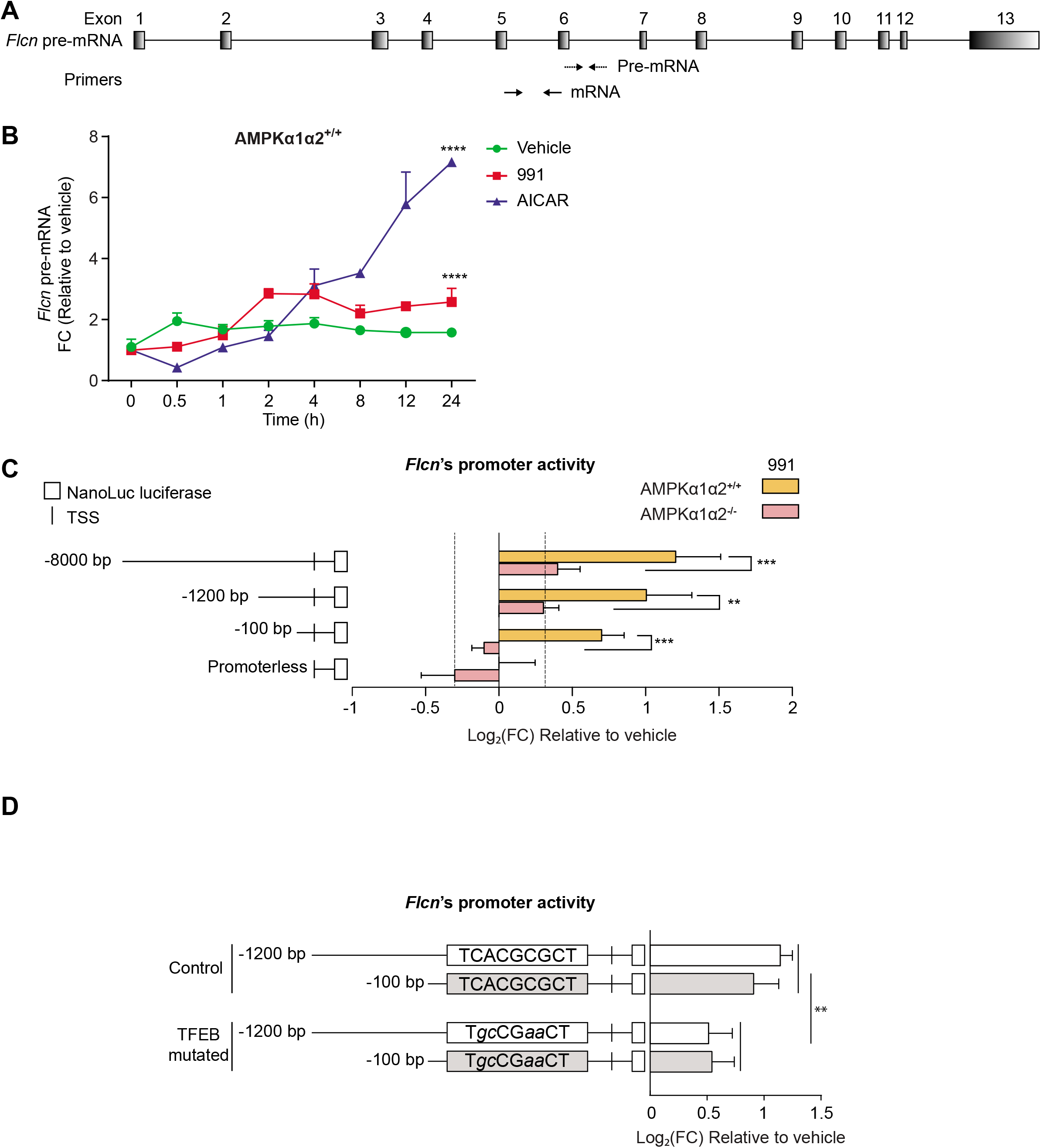
*Flcn* pre-mRNA and promoter activity. (**A**) Schematic representation of the primers designed to measure the pre-mRNA (exon-intron) and mRNA (exon-exon) of *Flcn* (**B**) Pre-mRNA level of *Flcn* in MEF AMPKα1α2^+/+^ treated with vehicle (DMSO), 10 μM 991 or 2 mM AICAR for 0, 0.5, 1, 2, 4, 8, 12 and 24 h. Data were analyzed by two-way ANOVA with the factors of time and treatment, plus the interaction between these two factors (n=3). Significance of the treatment factor is indicated, \*\*\*\**P* < 0.0001. (**C**) Schematic representation of *Flcn*-NanoLuc luciferase reporter plasmid and the correspondent luciferase activity profile. Several lengths of *Flcn*’s promoter region (−8000, −1200 and −100 bp) were inserted upstream of the NanoLuc luciferase reporter gene. The constructs, along with a firefly luciferase plasmid which served as internal control, were transiently transfected into MEF AMPKα1α2^+/+^ or AMPKα1α2^-/-^. Cells were then treated with vehicle (DMSO) or 30 μM 991. Values were first normalized by transfection efficiency, and then represented as log2 fold-change ± SD, relative to control (vehicle-treated cells) (n=3). The color corresponds to the three treatment conditions as indicated in the key. The grey shaded area indicates the log2 fold-change threshold of ±0.37. Data were analyzed by two-way ANOVA with the factors of genetic background and promoter length, plus the interaction between the two factors. Significance of the genetic background is indicated, \*\*\**P* < 0.001, \*\**P* < 0.01. (**D**) MEF WT were transiently transfected with *Flcn*-NanoLuc luciferase (−1200 or −100 bp) control or with a mutation on TFEB binding site (position −40 bp from the TSS), together with a firefly luciferase plasmid, and treated as above. Values were first normalized by transfection efficiency, and then represented as log2 fold-change ± SD, relative to control (vehicle-treated cells) (n=3). The color corresponds to the cellular genotype as indicated in the key. Data were analyzed by a two-way ANOVA with the factors of promoter length and TFEB binding site status (control or TFEB-mutated), plus the interaction between these two factors. Significance of the TFEB binding site status is indicated, \*\**P* < 0.01.

**Supplementary Table 1. Transcripts differentially regulated in response to 991 and AICAR.**

The excel file containing the complete list is available as a separate document.

**Supplementary Table 2.**
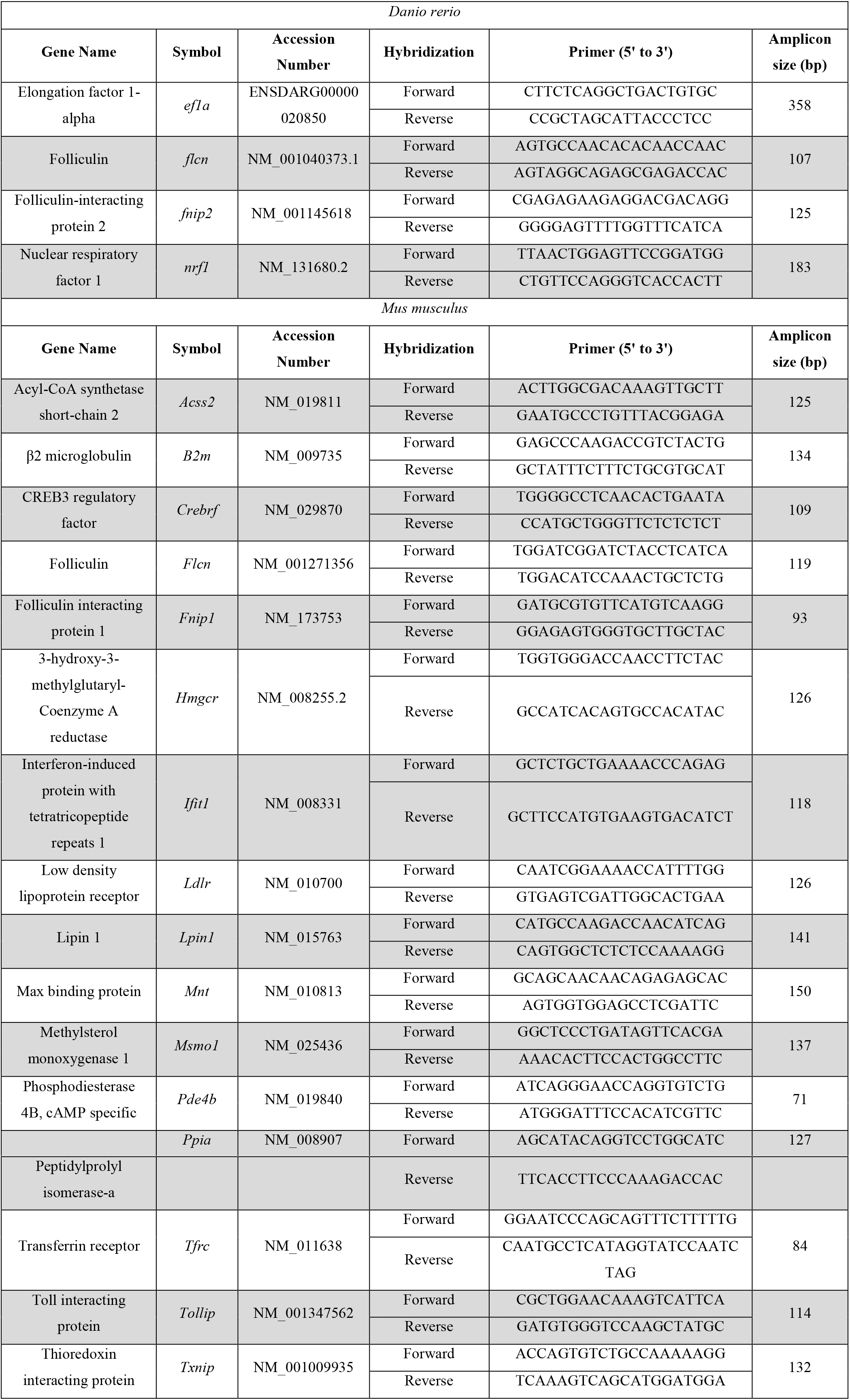
qPCR primers.

## Abbreviations

ACC: acetyl-CoA carboxylase
AICAR: 5-aminoimidazole-4-carboxamide ribonucleotide
AMPK: 5’-AMP-activated protein kinase
FLCN: Folliculin
TFEB: transcription factor EB

